# Evaluation of the Basis and Performance of Simple Allometry for Predictions of Clearance and Oral Clearance in Humans and Improvement Potential with Added *In Silico* Predictions and 2D PK-Space Positioning

**DOI:** 10.1101/2025.05.05.652242

**Authors:** Urban Fagerholm

## Abstract

**Background:** Allometry is traditionally used in drug development in order to extrapolate and predict pharmacokinetic (PK) parameter estimates, such as steady-state volume of distribution (V_ss_), clearance (CL), oral clearance (CL/F, where F is the oral bioavailability) in animal species to humans. Recent results show that *in silico* prediction methodology can improve and outperform laboratory data-based methods for predictions of V_ss_ and F. The main objectives were to evaluate the simple allometry principle for CL and CL/F, how well simple allometry predicts CL and CL/F in humans, and whether a combination of simple allometry and *in silico* predictions can improve predictions of CL and CL/F in humans.

**Methods:** The literature was searched for CL and CL/F-data in animal species and man. Data from at least 2 species, and preferably 3 or 4, and humans, for each compound (only small drugs) were used for the evaluation. The software ANDROMEDA by Prosilico was used for *in silico* predictions. A rule-based approach, based on laboratory and *in silico* data, and 2D PK-Space, were used to localize and predict compounds with largest allometric errors.

**Results and Discussion:** The evaluation shows limited support for the theoretical basis and empirical evidence and applicability of simple allometry for the prediction of CL and CL/F. There are many deviations from the simple allometric relationship (such as relatively low liver weight and blood flow in humans and 140-fold underprediction to 5,800-fold overprediction), skewness (including general overprediction, cases where humans deviate, and relatively high CL and low F in rats) and limited interspecies relationships (R^2^=0.07-0.19 for CL in animals *vs* humans). With 43 qualified compounds, R^2^ for CL and CL/F with simple allometry reached ca 1/3 (log scales). With the combination of simple allometry and *in silico* it was possible to improve the predictive performance. 31 to 124 % (average 68 %) improvement were found for R^2^, <2-fold error and median error, whereas maximum errors were reduced to 1/13 to 1/6. *In silico* predictions alone were even better - 19 to 176 % (average 96 %) and 1/157 to 1/25, respectively. 3 and 4 different PK-rules for large allometric prediction errors were found for CL and CL/F, respectively. These also had distinct positioning in 2D PK-Space. With *in silico* predictions, errors for the compounds with largest allometric prediction errors were decreased by 7-fold for CL and 4-fold for CL/F (median).

**Conclusion:** Simple allometry is associated with limited theoretical and empirical support for predictions of CL and CL/F in humans, and can be clearly improved when combined with (or replaced by) new *in silico* prediction methodology, rules and 2D PK-Space positioning. This is in line with the ambition to reduce and replace animal testing in drug development and need for methodological improvement.

## Introduction

Allometry is traditionally used in drug development in order to extrapolate and predict pharmacokinetic (PK) parameter estimates, such as steady-state volume of distribution (V_ss_), clearance (CL), apparent clearance following oral compound administration (CL/F, where F is the oral bioavailability) and systemic exposure (AUC_iv_=Dose/CL; AUC_po_=Dose•F/CL) from animal species to humans. Still, despite criticism [1-3], long known extreme maximum prediction errors (health risks) [2, 4, 5], and the fact that Dedrick, one of the pioneers in the field and famous for his work around allometry and PK-predictions, emphasized early on (in 1973) *“drug metabolism, in general, probably does not bear a relationship to body size”* [6], it appears to have general acceptance among drug developers and authorities. Every year between approximately 60 to 100 research articles based on allometric scaling are published [3].

For V_ss_, it has been shown by many that animal data are more predictive than *in vitro* data, and that addition of *in vitro* data can improve animal data-based predictions (on average ca 20-30 % improvement for a variety of statistical performance measures) [7-9]. The reduction of maximum errors is more extensive (several-fold) [7, 9]. Limitations with the inclusion of *in vitro* data, such as unbound fractions (f_u_) in plasma and tissues, include variability between and within laboratories (3.3-, 1.5- and 47-fold mean, median and maximum interlaboratory variability for f_u_ in human plasma, respectively [10]), cases with non-quantifiable estimates), and the opportunity to select favorable f_u_-values, and thereby manipulating the predictive performance [11].

In studies by, for example, Do Jones et al. (PhRMA CPCDC Initiative [7]); Petersson et al. [8] and Fagerholm et al. [12], *in silico*-based prediction methodology (by SimulationsPlus) was inferior to animal data-based methods. With our human clinical ADME/PK software ANDROMEDA by Prosilico [12, 13], we found that *in silico* methodology performed on par with simple allometry (based on data from rats, dogs and monkeys) and outperformed (head-to-head comparison; hold-out set for *in silico* predictions) *in vitro*+*in silico* methodology (SimulationsPlus) for 103 compounds [12]. A combination of simple allometry and *in silico* (ANDROMEDA) improved the performance with on average ca 30 % - R^2^ (0.74 *vs* 0.67 for log V_ss_), mean error (2.1- *vs* 2.6-fold), median error (1.4- *vs* 1.7-fold), maximum error (15- *vs* 29-fold) and %<2-fold error (70 *vs* 63 %) (Figure 1). The improvement is of same size as shown for animal data + *in vitro* data-based predictions, with the advantage that it does not have the limitations of laboratory methods (variability and non-quantifiable compounds; see above).

**Figure 1.**
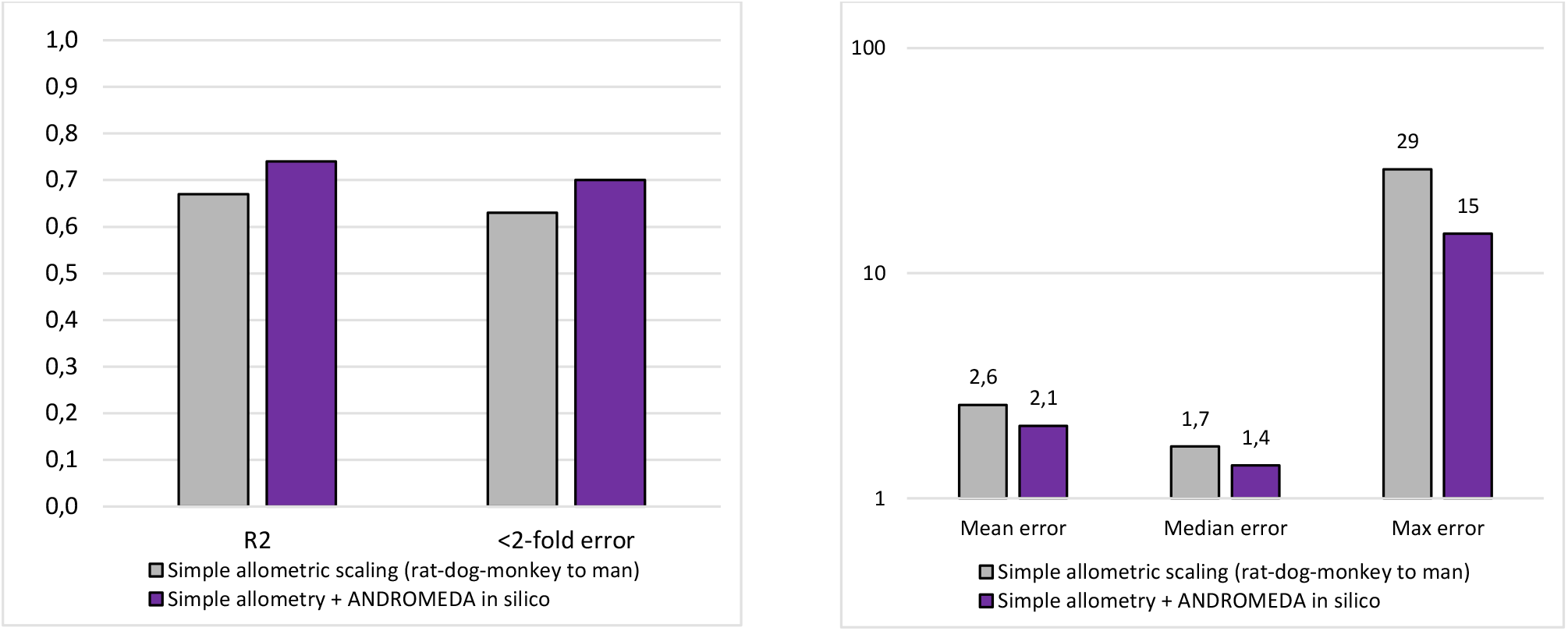
R^2^ and fraction of compounds with <2-fold errors (left), and mean, median and maximum errors (right), with simple allometry and a combination of simple allometry and *in silico* methodology (ANDROMEDA by Prosilico) for the prediction of V_ss_ in man.

**Figure 2.**
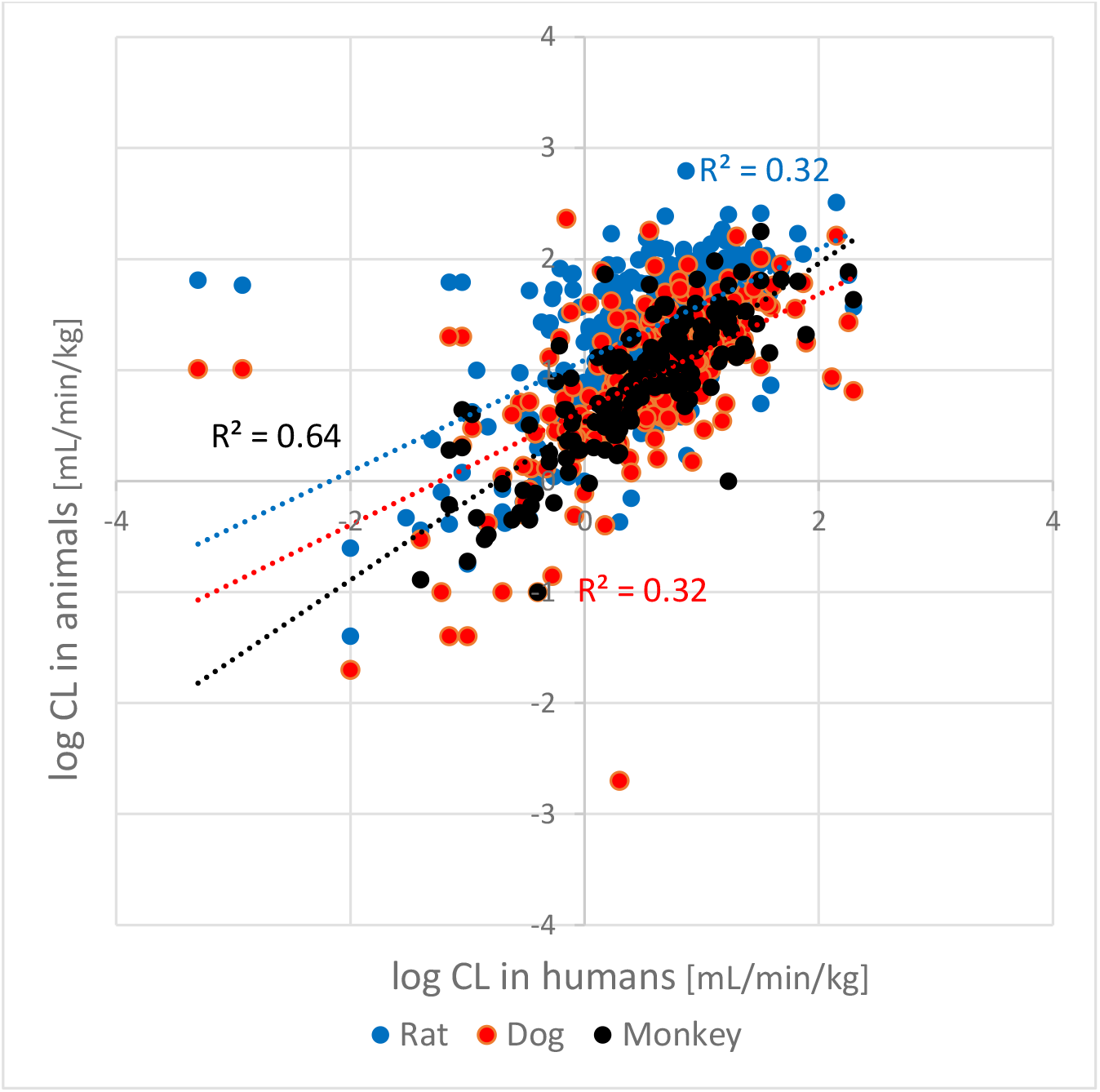
Relationships/correlations between rat, dog and monkey log CL *vs* human log CL for 309 compounds. R^2^-values for rat *vs* man, dog *vs* man and monkey *vs* man CL are 0.07 (n=296), 0.07 (n=245) and 0.19 (n=167), respectively. Corresponding R^2^-values for log CL are 0.32, 0.32 and 0.64, respectively.

Corresponding results obtained with ANDROMEDA alone were 0.65, 2.2-fold, 1.5-fold, 16-fold and 69 %, respectively. (Note: the *in silico* method underpredicted V_ss_ for compounds that bind to DNA and bone, but enables prediction of potential for enterohepatic recirculation, and thereby, higher V_dβ_ and longer half-life (t_½_)). These results show that *in silico* prediction methodology can both improve and outperform laboratory methods.

Allometric predictions of CL and CL/F (and AUC_iv_ and AUC_po_) in humans are not as straightforward and accurate as for V_ss_. The main objectives of this study were to evaluate 1) the simple allometry principle (theoretical and empirical basis) for CL and CL/F, 2) how well simple allometry predicts CL and CL/F in humans, and 3) whether a combination of simple allometry and *in silico* predictions (using ANDROMEDA by Prosilico and positioning in 2D PK-Space) can improve predictions of CL and CL/F in humans. A secondary aim was to evaluate average animal data, alone and in combination of *in silico* predictions, for the prediction of F in humans.

## Materials & Methods

The literature was searched for useful CL-, CL/F- and F-data in animal species and man. Data from at least 2 species, and preferably 3 or 4, for each compound (only small drugs) was set as a criterium. ANDROMEDA by Prosilico (version 2.0; major MW domain 150-750 g/mol) was used for corresponding (head-to-head) *in silico* predictions of human CL, F and CL/F. Results for the combination of animal and *in silico* data-based CL- and CL/F-predictions were calculated as the mean of logarithmic results from both methodologies. For F, a linear approach was applied and *in silico* prediction results were taken from a previous study [5].

A rule-based approach, based on laboratory data in animals and humans and *in silico* data (generated with ANDROMEDA), and 2D PK-Space (a 2-dimensional description of PK-Space in humans, based on clinical data for major PK-parameters of marketed small drugs and factor analysis; PK-Space positioning is based on observed or predicted PK-estimates; a 2-factor solution explains ca 80 % of the variance), were used to localize and predict compounds with largest allometric prediction errors.

Overall improvements of performance were estimated from the average improvements of R^2^, <2-fold error and median error, and from the ratio of maximum errors.

## Results

### Evaluation of the simple allometric principle for CL and CL/F

Simple allometric scaling of PK parameters is based on several assumptions (see Table 1). Apparently, there are many deviations from allometric relationships, skewness and limited interspecies relationships likely to create poor predictions for a significant number of compounds. Overall, the findings give limited support for simple allometry as a reliable prediction method for human CL and CL/F. This is in agreement with Van Valkengoed et al. [3], who also highlighted that many key assumptions are disputed or disproven and the increasing body of evidence against the existence of one universal allometric exponent.

**Table 1.** An evaluation of simple allometry principles (theoretical and empirical basis) for CL and CL/F.

| Assumptions | Findings |
| --- | --- |
| There is no or little deviation of blood flows (Q, mL/min) in major eliminating organs, such as the liver and kidneys, between species. | There is a simple allometric relationship for liver Q ($Q_H$), but the allometrically predicted $Q_H$ in humans is ca 50 % higher than the actual value (ca 2,150 vs 1,500 mL/min) [14]. This proposes a general overprediction potential of human CL and CL/F. |
| There is no or little deviation of weights (g) in major eliminating organs, such as the liver and kidneys, between species. | There is a simple allometric relationship for liver weight, but the allometrically predicted estimate in humans is roughly 20 % higher than the actual value [14]. This proposes a general overprediction potential of human CL and CL/F. |
| Active elimination processes, such as intrinsic hepatic metabolic and biliary elimination and active renal secretion, and plasma protein binding and reabsorption, show body weight dependency or are similar across species. | Figure 2 shows the quite poor relationships/correlations between rat, dog and monkey CL vs human CL for 309 compounds (data mainly taken from [9, 15]. $R^2$-values for rat vs man, dog vs man and monkey vs man CL are 0.07 (n=296), 0.07 (n=245) and 0.19 (n=167), respectively. Corresponding $R^2$-values for log CL are 0.32, 0.32 and 0.64, respectively (Figure 2). This is in line with [16], who found virtually no apparent relationship between intrinsic metabolic CL ($CL_{int}$) in rat and human microsomes for hundreds of compounds, and also what Dedrick, one of the pioneers in the field and famous for his work around allometry and PK-predictions, emphasized early on (in 1973) *“drug metabolism, in general, probably does not bear a relationship to body size”* [6]. Rat and mouse have relatively high CL and CL per kg body weight (compared to many animal species and man). For example, rats have ca 50 % higher median CL than predicted from simple allometry [16] and twice as high average CL/$Q_H$-ratio than humans (0.8 vs 0.4) [9, 15, 18]. Many compounds with high CL in rats (in total, 41 %) do not have high CL in man (in total, 14 %). For compounds with CL of at least 70 % of $Q_H$ (high liver extraction) in rats, dogs and monkeys, 46 % also have high CL in humans. Eighteen % of them have low human CL [9, 15, 18]. For compounds with low CL in these three animal species, 98 % also have low CL in humans. None of these are highly extracted by the human liver. The results indicate limited support (assumed limited predictive performance, with overprediction potential) for simple allometry for CL and CL/F. |
| First-pass processes such as dissolution, permeation, gut-wall extraction and liver extraction are | A dataset consisting of 78 compounds with F-values from at least 2 animal species and man [5, 19] showed irregular results, with 59 % of compounds that do not show allometric relationships, 18 % with allometric relationship, and 14 % that are difficult to classify. |
| body weight dependent or are similar across species. | $R^2$-estimates for F in mice, rats, dogs and monkeys vs humans are 0.25 (n=30), 0.29 (n=123), 0.38 (n=125) and 0.71 (n=40), respectively, with intercepts of 25-40 % human F. On average, humans have ca 40 % (relative %) higher F than in mice, rats and monkeys, and similar F as in dogs. In 7 % of cases, human F is lower than in 3 animal species, and in 23 % cases where F data are available from 1 or 2 animal species, human F is the lowest. Eleven compounds with F<15 % in all investigated animal species had F >50 % in man.<br>The findings show that F has limited/no allometric applicability. |
| If CL follows the simple allometric principle, this can also be assumed for CL/F. | This is most likely to occur in cases of low CL and high F in animals and humans. Since CL is a determinant for F, deviations from an allometric relationship are expected in cases of higher CL and lower F. This predicts that CL in humans is overall better predicted than CL/F using simple allometry.<br>In case human CL and CL/F are predicted using allometry and human F predicted in a different way one will end up having multiple and (probably) different CL and F-estimates to use. |

### Evaluation of the performance of simple allometric principle for CL

#### Published studies

An overall and clear systematic skewness (overpredictions) has been found for CL-predictions using simple allometry (using data from at least 3 animal species per compound) [18, 20]. This overprediction tendency is on average 1.7-fold (1.4-fold for median), which corresponds to almost 300 mL/min, and is most apparent and significant for drugs with low and high CL in humans [18]. Roughly 20 % of 61 investigated compounds were predicted to have a CL≥Q_H_ (seven compounds with predicted CL of 2,000-3,100 mL/min; consistent with zero F if assuming that CL=CL_H_) [18], and every other compound had an overprediction of more than 2-fold [20]. Thus, there is a potential risk with the simple allometric method to underestimate the potential of candidate drugs and overexpose humans in first clinical studies. The finding is consistent with the comparably low liver weight and blood flow in humans (see above).

These findings are in contrast to the suggestion by Zou et al. [21], who meant that simple allometry for CL is applicable for compounds with high hepatic extraction ratio (CL highly dependent on Q_H_) and has limited usefulness and reliability for compounds with low and moderate hepatic extraction ratios (*“simple allometry usually fails to predict the CL of such compounds”*).

For 39 % of 309 found/selected compounds, allometric relationships are found for CL per kg body weight (rat>monkey>dog>man), but for only 2 %, CL per kg body weight is similar (<2-fold interspecies difference) across species [data taken mainly from 4, 9, 15, 18]. Thus, based on CL per kg body weight, the simple allometric principle is violated for a majority of compounds.

In line with the limited interspecies relationships for CL, slope coefficients (*b*) for simple allometry vary considerably, ranging from -1.2 to 2.2 [2, 18]. For ca 10 % of compounds *b* exceeds 1 [2, 18]. A negative slope implies that CL in humans is much lower than in animals, especially in species with lower body weight.

Simple allometry is also associated with extensive individual maximum prediction errors - from 140-fold underprediction to 85-, 120- and 5,800-fold overpredictions (the CL of UCN-01 was overpredicted by a factor of 5,800) [4, 15, 18, 20]. The predicted and observed t_½_ of UCN-01 and ACD855 were half a day *vs* several months (ca 150-fold underprediction) and 5 *vs* 68 days (15-fold underprediction), respectively. Such predictions jeopardize the success and human risk potentials.

*In vitro* metabolic CL_int_- and f_u_-data from animals and humans are sometimes added to allometric predictions of CL. They could improve overall predictive performance by 10-25 % and reduce maximum error with about 2-fold in studies with many compounds [9]. Limitations with such approaches include cases with non-quantifiable CL_int_ (occurs for a significant number of compounds [10, 11]) and/or f_u_ [10], laboratory variability [10], and significant excretion, and the facts that CL is less dependent on CL_int_ and f_u_ at higher hepatic extraction ratios.

#### The present study

The literature search resulted in the finding of 65 compounds with CL-data in animals and humans and for which simple allometry is applicable for predictions [5, 9, 15, 18]. R^2^-estimates for predicted (from rat, dog and monkey data) *vs* observed log CL was 0.32 (Figure 3). The median, mean and maximum prediction errors were 1.8-, 96- and 5,800-fold, respectively, with 70 % overpredictions and 57 % of predictions with <2-fold error.

**Figure 3.**
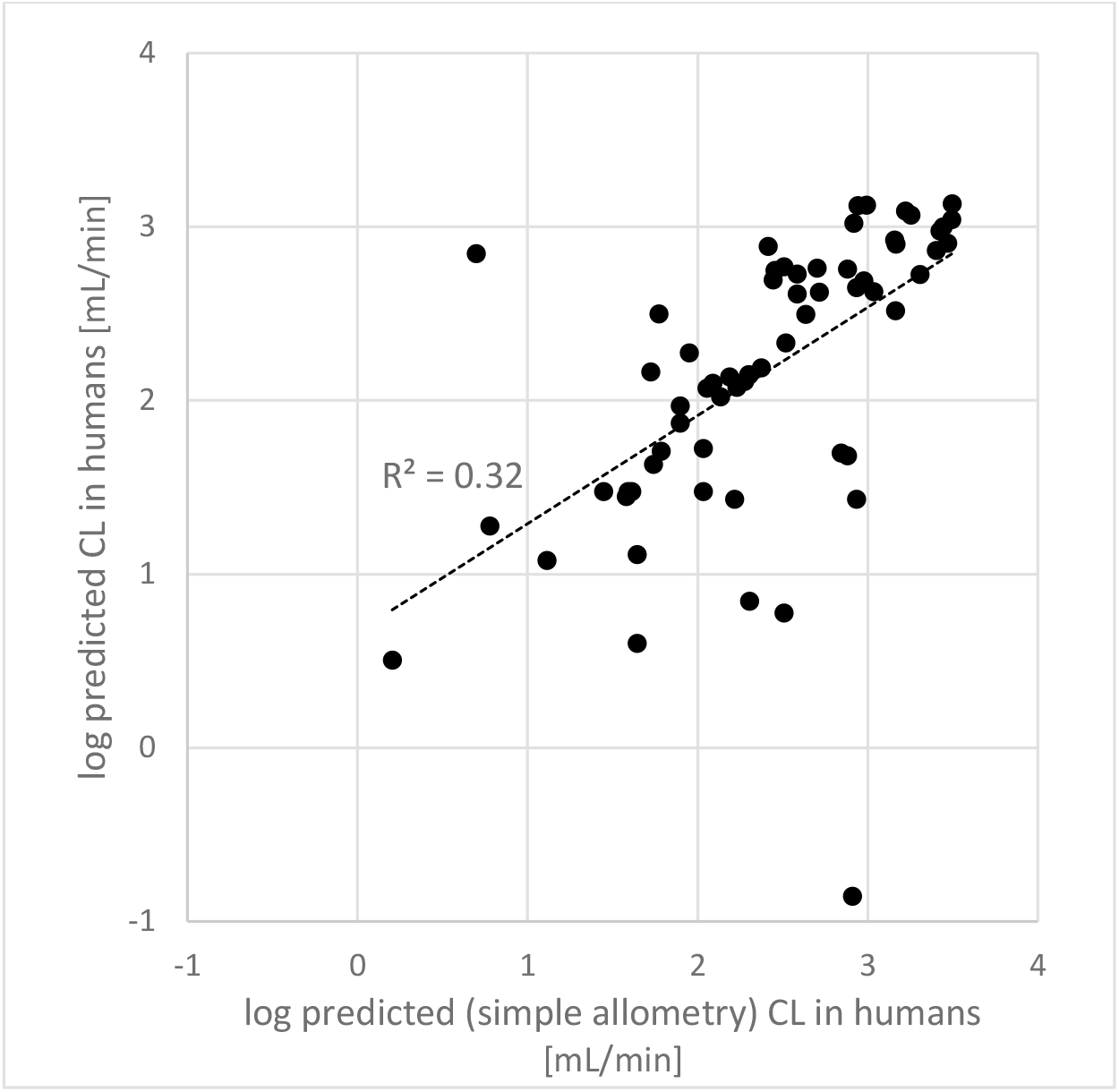
Predicted (simple allometry using data from rat, dog and monkey) *vs* observed log CL for 65 compounds.

Lombardo et al. [9] showed a similar number for simple allometry, 56 % of predictions with <2-fold error (n=97). They did, however, not include compounds with >100-fold prediction errors, such as UCN-01. Ring et al. [20] (PhRMA CPCDC Initiative) evaluated simple allometry for 18 compounds and reached R^2^=0.04 for log CL and 50 % of predictions with <2-fold error (overprediction trend).

### Evaluation of the performance of simple allometric principle for CL/F

#### Published studies

In a PhRMA CPCDC Initiative study [20], application of simple allometry for CL/F for 18 compounds resulted in under- and overpredictions of more than 100-fold, 34 and 71 % predictions with <2- and <10-fold errors, respectively, R^2^ of 0.03 (log scale) and the absolute average fold error of ca 5-fold. For the best PBPK-model used and tested in the PhRMA CPCDC Initiative 28 and 84 % of predictions reached <2- and <10-fold errors and the RMSE and absolute average error were ca 7.5- and 4.2-fold (n=82), respectively [22]. Predictions resulted in general overestimation of CL/F and underestimation of AUC_po_.

At AstraZeneca, laboratory data-based predictions of AUC_po_ for 58 internal compounds reached a R^2^ of 0.57 (log-scale) and RMSE of 0.59 (ca 4-fold) [23]. Corresponding results for their *in silico* prediction method were 0.33 and 0.74 (ca 5.5-fold), respectively.

#### The present study

Forty-three compounds with CL/F in at least 2 animal species (rat, dog and monkey; 17 compounds had measured CL/F in these 3 species) and in humans, and with MW between 150 and 750 g/mol, were found in the literature and selected for simple allometric predictions of CL/F in man [5, 9, 15, 18]. This selection includes azithromycin, erythromycin, acyclovir, amlodipine, ethinyl estradiol, etoposide, felodipine, fleroxacin, fluconazole, fluvastatin, furosemide, gabapentin, gatifloxacin, ketorolac, lidocaine, melagatran, midazolam, morphine, oseltamivir, propranolol, quinidine, remoxipride, sumatriptan, theophylline, venlafaxine, verapamil and UCN-01. *b* for all except two were positive (range -1.4 to 1.9). Mean, median and maximum errors were 51-, 4.7- and 1,790-fold, respectively (tacrolimus, with a MW of 804 g/mol and 4717-fold overprediction, was excluded). The R^2^ for log predicted *vs* observed CL/F was 0.36 (Figure 4; 0.32 when tacrolimus was included), with overprediction trend at low CL/F and underprediction trend at high CL/F. 21 and 70 % of predictions had errors <2- and <10-fold, respectively.

**Figure 4.**
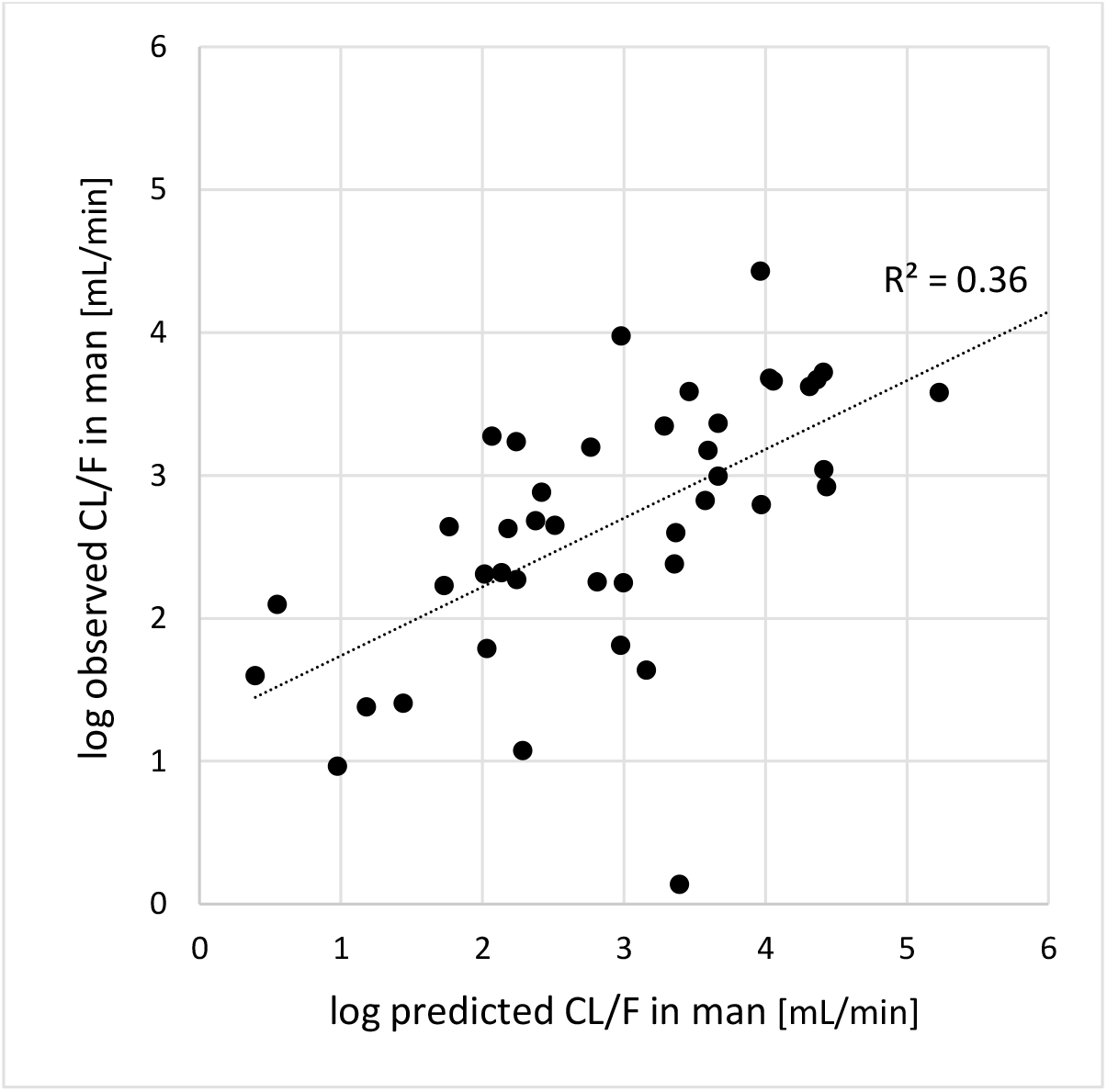
Predicted (simple allometry using data from rat, dog and monkey) *vs* observed log CL/F for 43 compounds.

For this set of compounds MW, fraction renally excreted (f_e_), f_u_, CL and F ranged from 171 to 749 g/mol, 0 to 1 (12 compounds with f_e_>0.5), <0.0002 to 0.97, 8 to 1,680 mL/min, and 4 to 100 %, respectively. About every fourth compound are known and/or predicted to be excreted unchanged via bile. However, only very few have limited passive permeability (1) or dissolution limitation (4) (and potentially very low F and higher uncertainty for CL/F-predictions), and no compound has very low passive permeability and very limited dissolution potential.

### Evaluation of the addition of *in silico* predictions

#### CL/F

The combination of simple allometry and *in silico* methodology (ANDROMEDA) for the 43 selected compounds improved the predictive performance compared to simple allometry alone (Figures 5 and 6). The R^2^ for log predicted *vs* measured CL/F with the combination of methodologies was 0.52 (*vs* 0.36 with simple allometry) and the systematic error trend was markedly reduced compared to simple allometry alone. Mean, median and maximum errors were of 11-, 2.6- and 310-fold, respectively. Thus, prediction errors were reduced by ca 5-, 1.8- and 6-fold, respectively. 47 and 93 % of predictions had errors <2- and <10-fold, respectively, which is also markedly better than with simple allometry alone (21 and 70 %, respectively).

**Figure 5.**
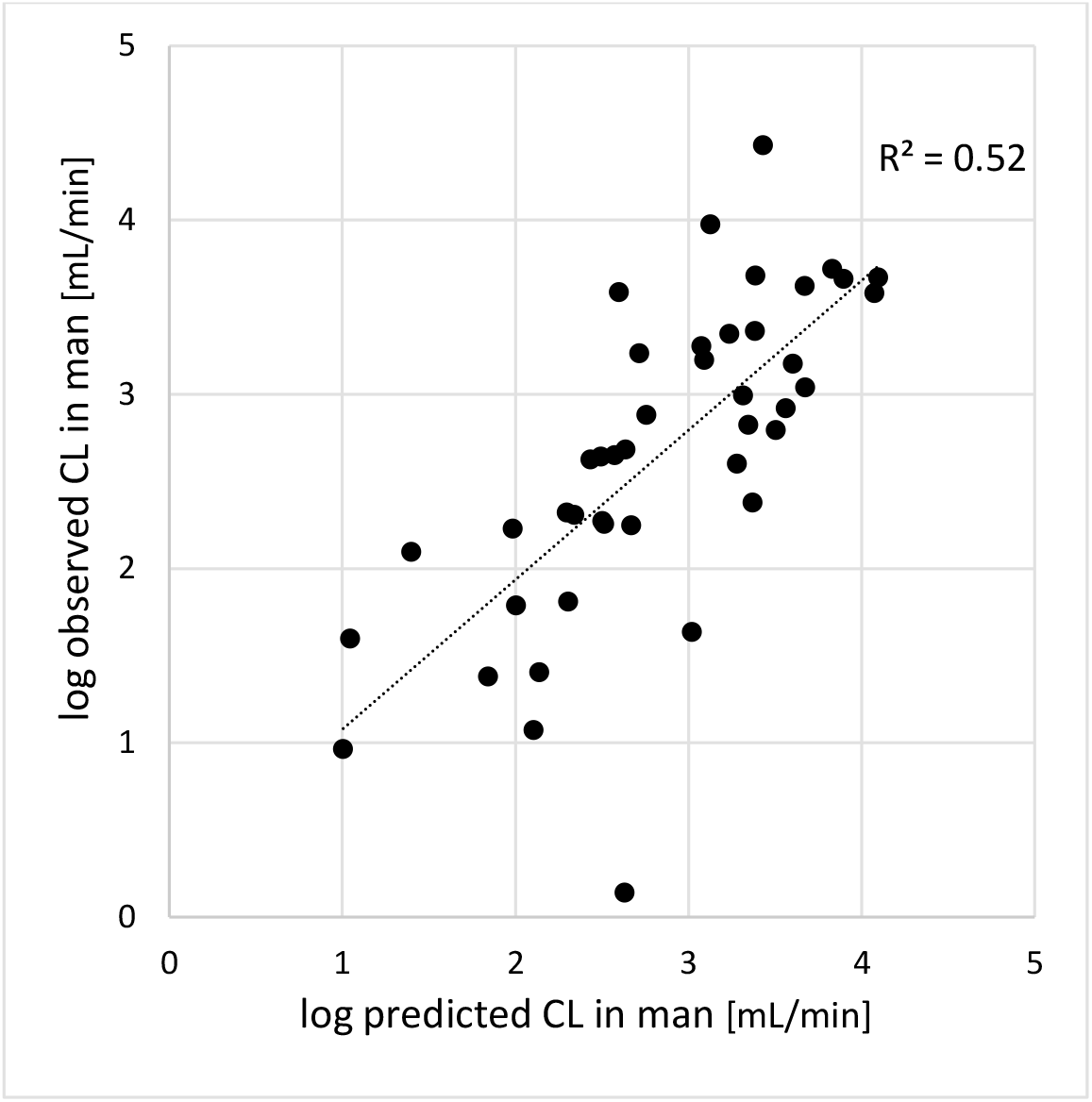
Predicted (combination of simple allometry using data from rat, dog and monkey + ANDROMEDA *in silico* methodology) *vs* observed log CL/F for 43 compounds.

**Figure 6.**
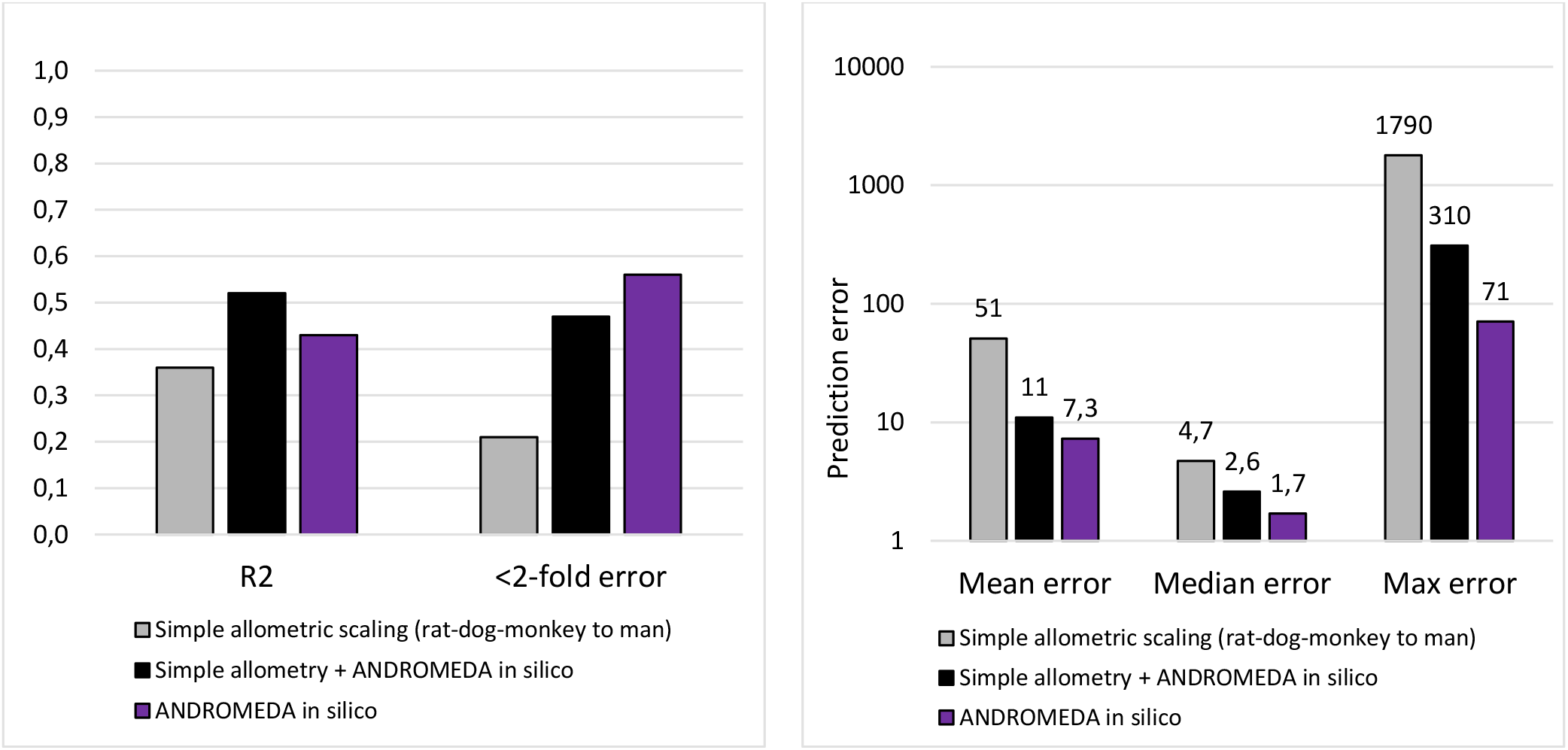
R^2^ and fraction of compounds with <2-fold errors (left), and mean, median and maximum errors (right), with simple allometry, a combination of simple allometry and *in silico* methodology (ANDROMEDA) and ANDROMEDA alone for the prediction of CL/F for 43 compounds in man.

The *in silico* methodology (ANDROMEDA) alone produced the following results for CL/F - Q^2^=0.43, 56 % with <2-fold error, 86 % with <10-fold error, 7.3-, 1.7- and 71-fold mean, median and maximum errors, respectively. Thus, it performed overall better than simple allometry and the combination of the two methodologies.

#### CL

The combination of simple allometry and *in silico* methodology (ANDROMEDA) also improved the predictive performance *vs* simple allometry for CL (same 43 compounds as used for CL/F evaluation above) (Figure 7). The R^2^ for log predicted *vs* measured CL with the combination of methodologies was 0.37 (*vs* 0.20 with simple allometry) and the systematic error trend was markedly reduced compared to simple allometry alone for this parameter. Mean, median and maximum errors were of 13-, 1.6- and 454-fold (454-fold for UCN-01), respectively. Corresponding estimates for simple allometry were 139- (11 times higher than the combination), 2.1- (1.3-fold higher) and 5800-fold (13-fold higher), respectively. 67 and 91 % of predictions had errors <2- and <10-fold, respectively, which is also better than with simple allometry alone (47 and 88 %, respectively).

**Figure 7.**
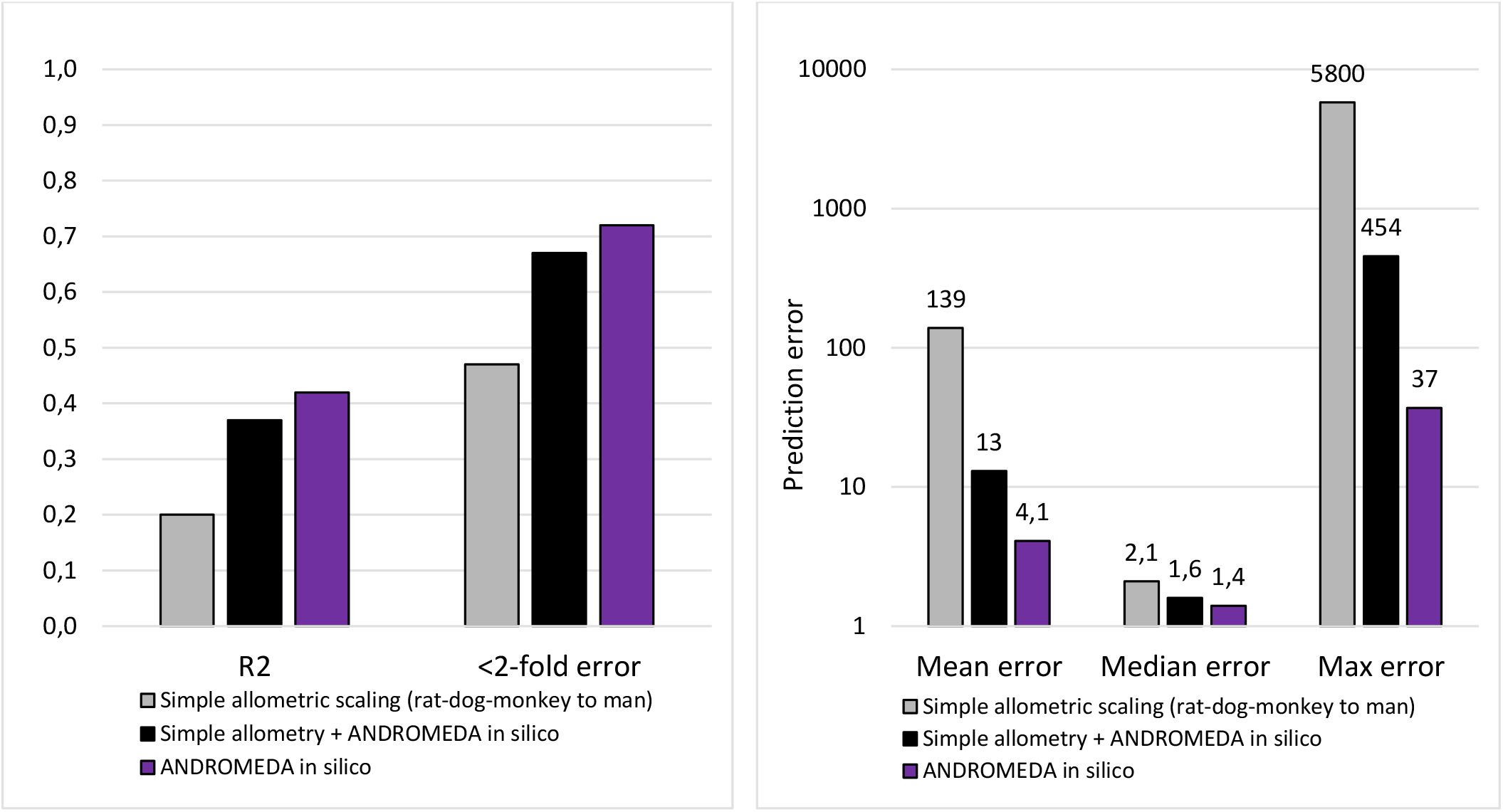
R^2^ and fraction of compounds with <2-fold errors (left), and mean, median and maximum errors (right), with simple allometry, a combination of simple allometry and *in silico* methodology (ANDROMEDA) and ANDROMEDA alone for the prediction of CL for 43 compounds in man.

The *in silico* methodology (ANDROMEDA) alone produced the following results for CL - Q^2^=0.42, 72 % with <2-fold error, 95 % with <10-fold error, 4.1-, 1.4- and 37-fold (36-fold overprediction of CL for UCN-01, which is known for its unusual PK and poor predictions and specifically high binding to α_1_-acid glycoprotein [5]) mean, median and maximum errors, respectively. Thus, and consistent with CL/F-predictions, it performed overall better than simple allometry and the combination of the two methodologies.

Within major pharmaceutical companies Q^2^s and RMSEs for *in silico* CL-prediction methods have reached 0.05 (log scale; n=343) [24] to 0.1-0.2 [23] and 0.4 to 0.7 (ca 2.5-to 4.5-fold) [20, 22, 23], which is inferior to results obtained with simple allometry.

#### F

Using a dataset consisting on F-values for 78 compounds in at least 2 animal species and man [5, 19] showed irregular results violating the allometric principle. 59 % of the compounds did not demonstrate allometric relationships, whereas 18 % did and 14 % were difficult to classify.

R^2^-estimates for mouse, rat, dog and monkey *vs* man were 0.25 (n=30), 0.29 (n=123), 0.38 (n=125) and 0.71 (n=40), respectively, and the intercepts were 25-40 % human F.

On average, and based on available data, humans have ca 40 % (relative %) higher F than in mice, rats and monkeys and similar F as in dogs. In 7 % of cases, human F is lower than in 3 animal species, and in 23 % cases where F data is available from 1 or 2 animal species, human F is lowest. For 11 compounds with F <15 % in all investigated animal species human F exceed 50 %.

When using the average F in at least two animal species to predict F in humans, R^2^ was 0.46, 65 % of predictions had <2-fold error and mean, median and maximum errors were 2.5-, 1.4- and 35-fold, respectively (n=78) (Figure 8). A combination of animal data and *in silico* data for F-predictions improved the outcome - R^2^=0.61, 74 % with <2-fold error and 2.1-, 1.3- and 23-fold mean, median and maximum prediction errors, respectively.

**Figure 8.**
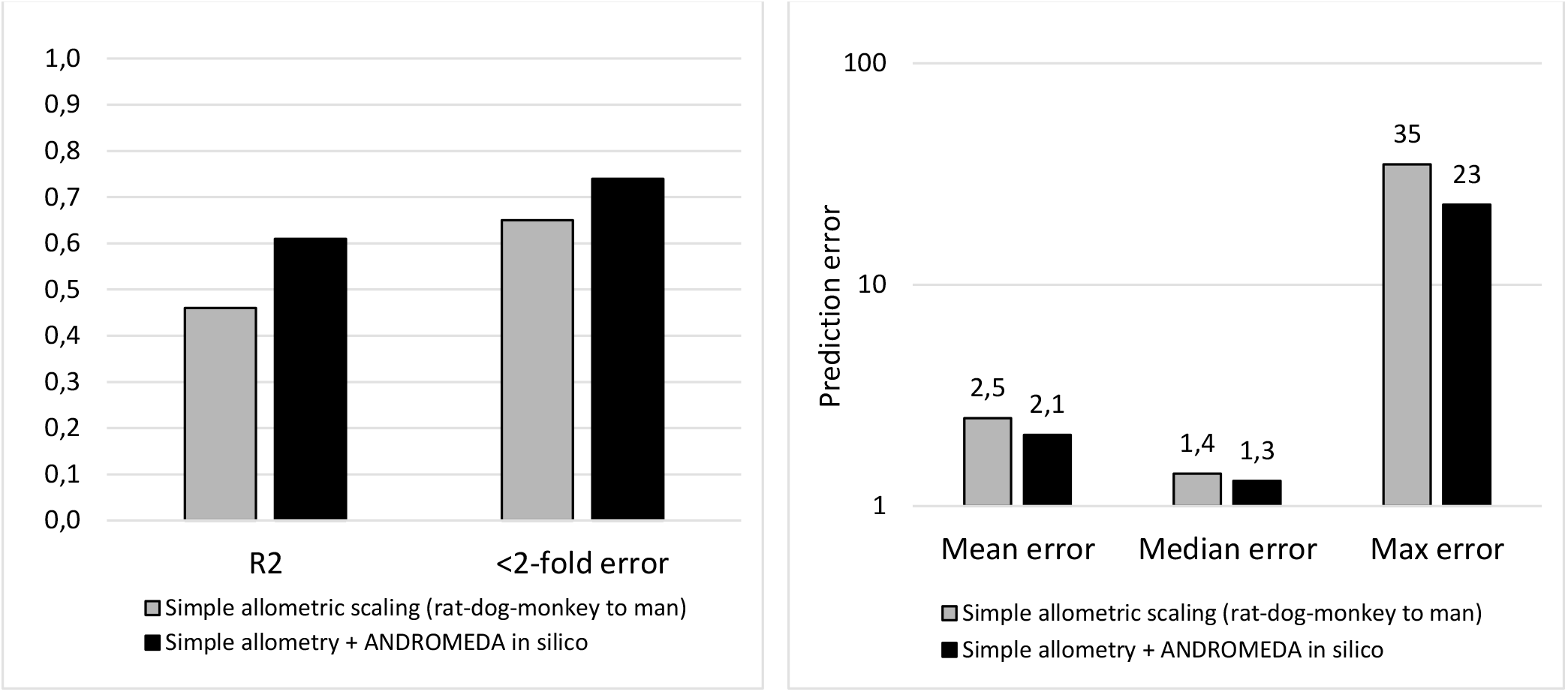
R^2^ and fraction of compounds with <2-fold errors (left), and mean, median and maximum errors (right), with simple allometry and a combination of simple allometry and *in silico* methodology for the prediction of F of 78 compounds in man.

### Rules and PK-Space for localization and improved predictions of compounds with large error

#### CL

For 14 compounds with largest allometric prediction errors of CL (4-to 5,800-fold; median 10-fold) a 3-rule-based system was found (all with different set-ups of PK-characteristics). These also had distinct positioning in 2D PK-Space (Figure 9). Compounds with potentially high allometric errors could be predicted via both the 3 rules and 2D PK-Space localization. 2D PK-Space zones for large and small predictions had approximately 50 % overlap, and about half of the 2D PK-Space was unexplored in this analysis. ANDROMEDA produced 7-fold lower median errors than simple allometry for these 14 compounds, and all except one (1.4-fold difference in favor of allometry) had higher prediction error with allometry than with ANDROMEDA.

**Figure 9.**
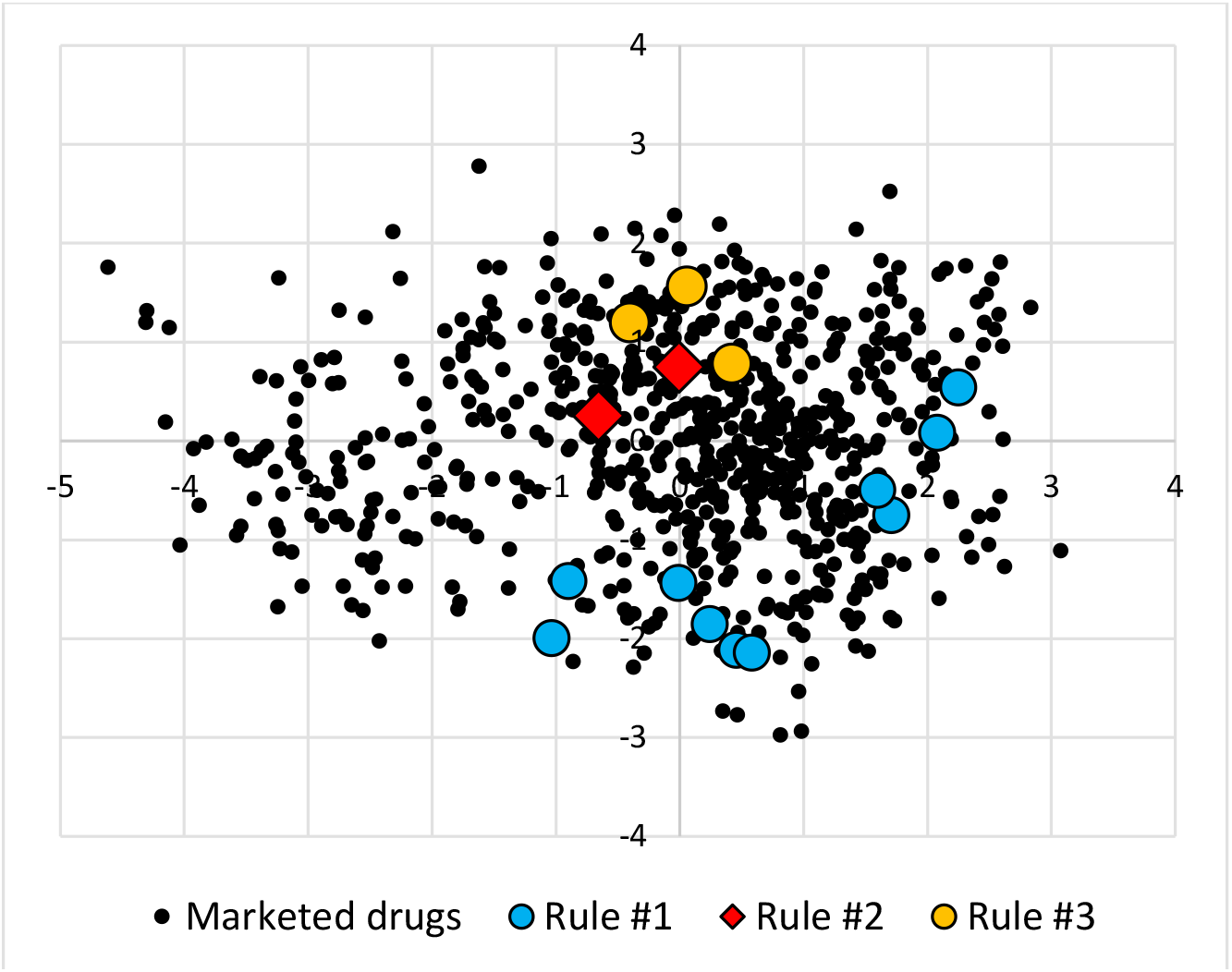
2D PK-Space (x=factor 1; y=factor 2; produced using factor analysis) positioning of 14 compounds with largest CL-prediction errors. The 14 compounds belong to 3 domains of compounds with certain PK-characteristics (Rules #1-3).

#### CL/F

19 of the 43 selected compounds had median and maximum allometric CL/F-prediction errors of 15- and 1,790-fold, respectively. Ten of these were also in the 3 groups of compounds with large errors for CL. An additional fourth rule was found for CL/F and the 9 compounds not classifying for rules #1-3. These had apparent distinct (but less clear as for CL) positioning in 2D PK-Space (Figure 10). ANDROMEDA produced 4-fold lower median errors than simple allometry for these 19 compounds, and all except one (1.06-fold difference in favor of allometry) had higher prediction error with allometry than with ANDROMEDA.

**Figure 10.**
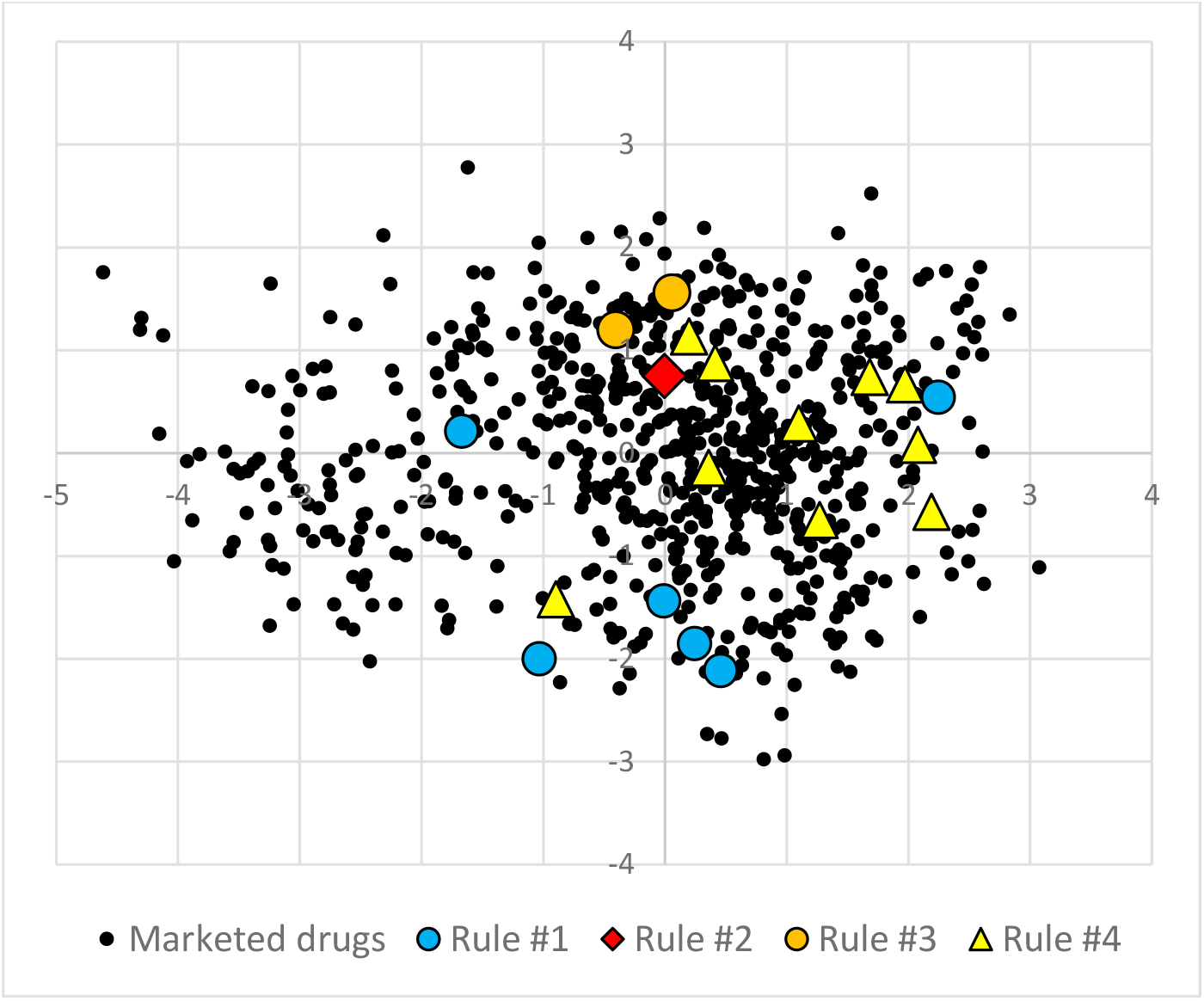
2D PK-Space positioning of 19 compounds with largest CL/F-prediction errors. The 19 compounds belong to 4 domains of compounds with certain PK-characteristics (Rules #1-3 as for CL and Rule #4).

## Discussion & Conclusion

This evaluation shows limited support for theoretical basis and empirical evidence and applicability of simple allometry for the prediction of CL and CL/F, and no clear support for the applicability of allometry for predictions of F in humans.

There are many deviations from simple allometric relationship (such as relatively low liver weight and Q in humans and 140-fold underprediction to 5,800-fold overprediction of CL), skewness (including general overprediction, cases where humans deviate, and relatively high CL and low F in rats) and limited interspecies relationships (R^2^=0.07-0.19 for CL in animal species *vs* humans). Such factors are never, or at least rarely, considered when producing confidence estimates for allometrically scaled human PK-

R^2^-values for CL and CL/F with simple allometry in the present study are ca 1/3 (log scales), which shows moderate predictive value with this methodology.

Addition of *in vitro* data to allometric scaling can improve the predictive performance of ca 10-30 % for various statistical measures and reduce maximum errors by 2- to several-fold (both for V_ss_ and CL). Limitations with such additions include laboratory variability (which often is several-fold), inability to quantify metabolic CL_int_ (common with microsomes and hepatocytes, but also for f_u_), and lower impact of f_u_ and CL_int_ at higher liver extraction ratios and when excretion occurs.

Inclusion/addition of data obtained in monkeys generally improves allometric predictions, but there are cases (at least 7 found) where CL in monkeys deviate markedly (several-fold) from other animal species and man and could lead to poorer human predictions (for example, for lidocaine). It is not recommended to use primates (for ethical reasons) for the generation of data for predictions of PK in humans (although this still seems to occur).

With the combination of simple allometry and ANDROMEDA (*in silico*) it is possible to improve the predictive performance and reduce maximum prediction errors for both CL, CL/F, and V_ss_. For CL, the improvement for R^2^ and <2-fold error and median errors, and for maximum errors, is 31-85 % (average 53 %) and 13-fold, respectively. Corresponding improvement numbers for CL/F are 44-124 % (average 83 %) and 5.8-fold, respectively. For F (*vs* average of animal F-values) the estimates are 8-33 % and 1.5-fold, respectively. For V_ss_, the improvements are 10-25 % and 2-fold, respectively [12].

*In silico* predictions alone were even better than the combinations of *in silico* and simple allometry – 50-110 % (average 71 %) and 157-fold for CL; 19-176 % (average 121 %) and 25-fold for CL/F. Another advantage with ANDROMEDA is that it produces estimates for many different ADME/PK-processes (including renal and biliary excretion and gut-wall extraction), and thereby, predicted descriptions of ADME/PK-characteristics for compounds.

The 4 different PK-rules, supported by 2D PK-Space positioning, for large allometric prediction errors are useful for finding and predicting compounds whose CL and/or CL/F are most likely to be high. ANDROMEDA produced markedly lower errors for all except one of these large error compounds, and is, therefore, useful for sanity-checking of rejection of available laboratory PK-data. See Figure 11 for how this approach can be used to localize and improve predictions for such compounds.

**Figure 11.**
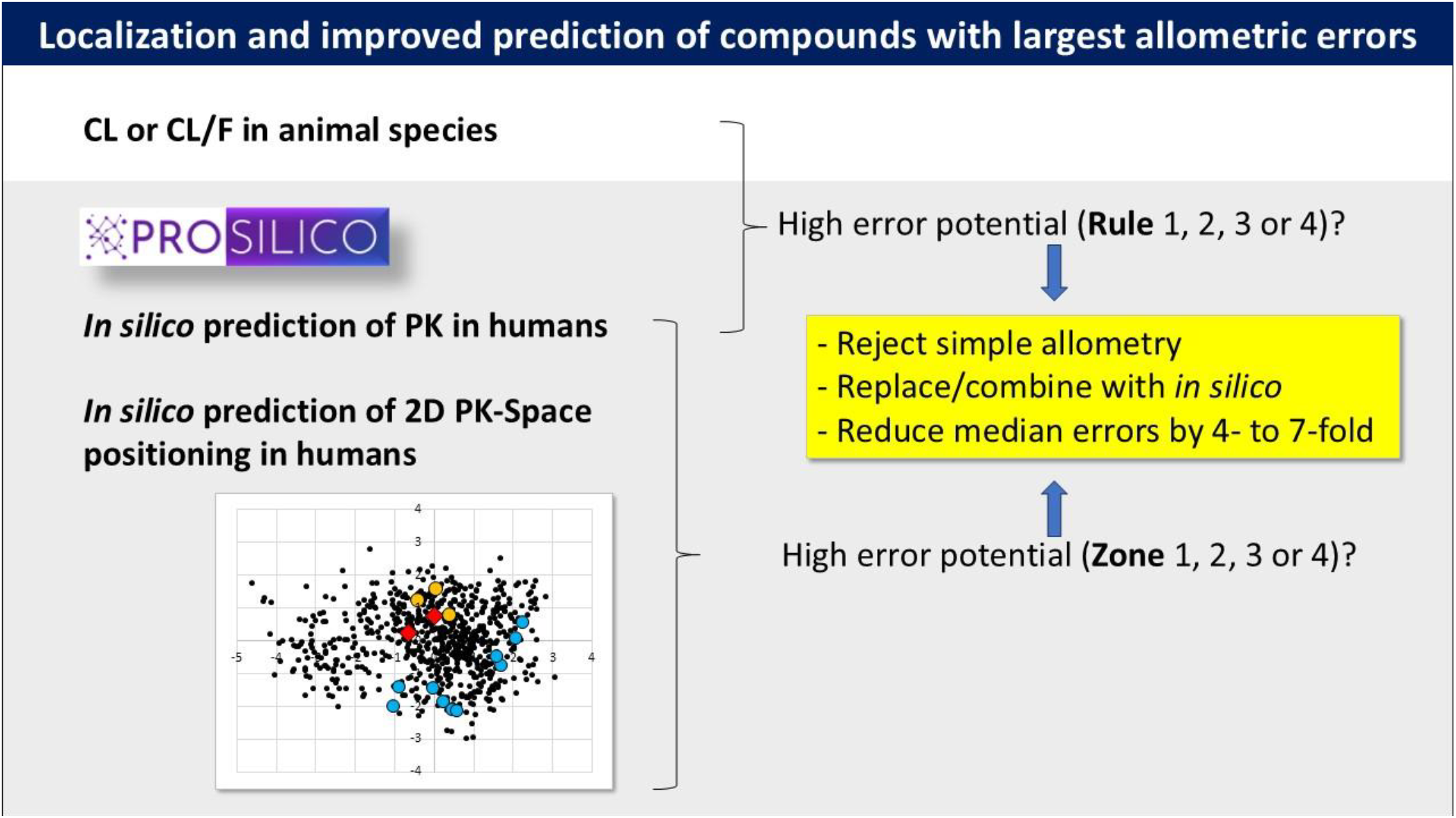
Flow chart for localization and improved CL- and CL/F-predictions of compounds with large allometric prediction errors.

Thus, it is possible to increase the accuracy and confidence and reduce the maximum errors, risks and use of animals with the use of ANDROMEDA, either alone or combined with laboratory data and models. These results and advances are in line with the important step towards the goal to reduce, refine and replace animal studies in drug discovery and development recently taken by FDA (*FDA Announces Plan to Phase Out Animal Testing Requirement for Monoclonal Antibodies and Other Drugs*) [25].

